# DNA Denaturing through Photon Dissipation: A Possible Route to Archean Non-enzymatic Replication

**DOI:** 10.1101/009126

**Authors:** Karo Michaelian, Norberto Santillán Padilla

**Affiliations:** Instituto de Física, UNAM. Cto. Interior de la Investigación Científica, Ciudad Universitaria, México D.F., C.P. 04510; Facultad de Ciencias, UNAM. Cto. Interior de la Investigación Científica, Ciudad Universitaria, México D.F., C.P. 04510

## Abstract

Formidable difficulties arise when attempting to explain the non-enzymatic replication, proliferation, and the acquisition of homochirality and information content, of RNA and DNA at the beginnings of life. However, new light can be shed on these problems by viewing the origin of life as a non-equilibrium thermodynamic process in which RNA, DNA and other fundamental molecules of life arose as structures to dissipate the prevailing solar spectrum. Here we present experimental results which demonstrate that the absorption and dissipation of UV-C light by DNA at temperatures below their melting temperature leads to complete and reversible denaturing for small synthetic DNA of 25 base pairs (bp), and to partial and reversible denaturing for 48 bp DNA and for large salmon sperm and yeast DNA of average size 100 kbp. This result has direct bearing on the above mentioned problems and thereby opens the door to a possible thermodynamic route to the origin of life.

## Introduction

Many of the fundamental molecules of life, those common to all three domains; bacteria, eukaryote, and archea, including RNA and DNA, amino acids, enzymes, vitamins, cofactors, and protoporphyrins, absorb photons in the UV-C^1^. RNA or DNA in complexes with these molecules act as acceptor quenchers, providing the electronically excited pigment donor molecule with an extremely rapid (sub picosecond) non-radiative dexcitation channel, through internal conversion into vibrational energy of the nucleic acid and surrounding water molecules^2^.

These pigment molecules have conjugated carbon bonds which not only gives them their UV-C absorption properties but also their planer aromatic-like structures and thus chemical affinity to RNA and DNA by intercalating between consecutive base pairs. Many also have chemical affinity to the secondary or tertiary structures of RNA and DNA^3^. The aromatic amino acids (tryptophan, tyrosine, phenylalanine), as well as two other UV-C absorbing amino acids (histidine and cystine), have particularly strong chemical affinity to their DNA codons or anti-codons^4^. This hints at, not only a stereochemical era,^4,5,6^ but also a UV-C dissipative era for early life,^7,8^ Sagan^9^ and Cnossen et al.^10^ have demonstrated the existence of an Archean atmospheric window covering precisely the absorption wavelengths of these pigments^1^.

Because these pigment-RNA/DNA complexes could have dissipated the Archean UV-C solar photon potential, their production and proliferation to well beyond expected equilibrium concentrations, before the appearance of biological metabolic pathways, could be explained as a non-equilibrium thermodynamic process which arose spontaneously to dissipate this photon potential^11^. This thermodynamic explanation of proliferation is applicable even today at visible wavelengths, irrespective of the complex contemporary metabolic pathways to the production of pigments^12^.

It has been conjectured that the origin and evolution of life was contingent on increases in the entropy production of the biosphere through increases in the dissipation of photons in the prevailing solar spectrum at Earth’s surface^7,8,12,13^. In particular, that RNA, DNA and the other fundamental molecules of life evolved under non-equilibrium thermodynamic directives to optimize the absorption and dissipation within the wavelength region of 230 to 290 nm, corresponding to a window in the Archean atmosphere^1,9,10^. The resulting heat from the dissipation of these photons could have denatured RNA and DNA, once the ocean surface temperature descended below the relevant melting temperature, without the need for enzymes, allowing for subsequent extension during overnight dark periods facilitated by Mg^2+^ ions^14^ and UV-activated phosphorylated nucleotides^15^. This proposed ultraviolet and temperature assisted reproduction (UVTAR) is similar to polymerase chain reaction (PCR)^16^ but in which the heating and cooling cycle is partially substituted by a day/night UV-C light cycling, and in which Mg^2+^ ions, or their complexes with a hypothetical molecule, played the role of the extension enzyme polymerase.

The UVTAR mechanism^7,8^ provides a plausible explanation of the homochirality of life^17^ as a result of the positive circular dichroism of RNA and DNA around 260 nm which corresponds to the peak in the UV-C solar spectrum reaching Earth’s surface during the Archean^1^, and a small prevalence of right over left handed circularly polarized submarine light in the late afternoon^18,19^ when surface water temperatures were highest and thus most conducive to denaturing. The UVTAR mechanism also provides a possible explanation for the beginnings of information accumulation in RNA and DNA^7,8^ since the aromatic amino acids that absorb in the UV-C have known chemical affinity to their codons or anti-codons^4,20^. RNA or DNA coding for these “antenna” amino acids would experience greater local heating under UV-C light dissipation and thus would enjoy greater reproductive success through the UVTAR mechanism in increasingly colder seas. Proliferation, leading to greater dissipation, would be thermodynamically driven^7,8^.

In this article we present unequivocal evidence for the reversible UV-C light denaturing of DNA for four different DNA samples; short 25 bp and 48 bp synthetic DNA, and relatively large (∼100 kbp) salmon sperm and yeast DNA. A quantitative analysis of hyperchromism and scattering in our extinction data suggests that UV-C light induced denaturing is significant and sufficiently large to support the explanations for homochirality and the accumulation of information in DNA through the dissipation based mechanisms mentioned above.

## Method

Salmon sperm and yeast DNA samples of varied lengths (average 100 kbp) were obtained from the Institute of Cellular Physiology at the National Autonomous University of Mexico (IFC-UNAM). Complementary synthetic DNA oligonucleotides of 25 and 48 bases were synthesized at the IFC-UNAM. The oligos were designed to be free of adjacent thymine and to have convenient denaturing and priming temperatures in order to facilitate a possible UVTAR mechanism operating under only very small simulated diurnal temperature cycling.

To form the double helices of 25 and 48 bp, complementary oligos at equal concentration were mixed in a Dulbecco PBS buffer (pH 7.3) solution containing 2.7 mM potassium chloride (KCl), 136.9 mM sodium chloride (NaCl), 1.5 mM potassium phosphate monobasic (KH_2_PO_4_) and 8.9 mM sodium phosphate dibasic (Na_2_HPO_4_), and heated rapidly to 85° C and kept there during 10 min before being brought to ambient temperature at a rate of 1° C/min.

The yeast DNA was dissolved in a Dulbecco PBS buffer and the salmon sperm DNA dissolved in purified water (Mili-q). The resulting concentrations of double helix DNA were determined from their absorption at 260 nm to be 2.2, 0.7, 0.0015 and 0.00023 μM for the 25 bp, 48 bp, yeast and salmon sperm DNA respectively (assuming average lengths of 100 kpb for the latter two).

One ml of the corresponding solution was placed in a standard quartz cuvette of 1 cm light path length. The cuvette was placed inside a precise (± 0.01°C) Ocean Optics^@^ temperature control unit operating via the Peltier effect with water flow stabilization of the temperature; important for precise temperature control when removal of heat from the cuvette was required. The temperature was monitored via a probe located in one of the four supporting towers of the cuvette and in direct contact with it. A magnetic stirrer provided assurance of temperature equilibration throughout the sample and homogeneous passage of the DNA through the approximately 3.7 mm diameter light beam on the sample.

UV light from a 3.8 W (9.4 μW on sample) deuterium source, covering continuously the range 200 to 800 nm, or light from a 1.2 W (7.0 μW on sample) tungsten-halogen source (used as a control) covering 550 to 1000 nm, was defocused onto the DNA sample via UV light resistant 30 cm long optical fiber of 600 μm diameter and a quartz lens. After passing through the sample, the surviving light was collected by a second lens and focused onto a similar optical fiber which fed into an Ocean Optics^@^ HR4000CG charge-coupled device spectrometer. The spectrometer covered the range 200 to 1200 nm with a resolution of 0.3 nm. Spectrometer integration times were of the order of 1.5 s and the data were averaged over five wavelength channels (1.5 nm).

After allowing approximately 1/2 hour for lamp stabilization, sample extinction (absorption plus scattering) spectra, *E*(λ), were obtained by comparing the measured intensity spectrum, *I*(λ), of the light through the DNA sample with a reference spectrum, *I_R_*(λ), obtained at room temperature with a similar quartz cuvette containing either purified water or PBS buffer as required, but no DNA. A dark spectrum, *I_D_*(λ), obtained with a shutter blocking the deuterium light to the reference sample at room temperature, was subtracted from the spectra to correct for stray light and electronic noise. Extinction was thus calculated as;

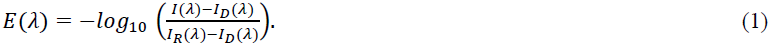

The synthetic DNA samples were heated rapidly (5°C/min) to 90 ° and held there for approximately 10 minutes to ensure complete denaturing. The temperature of the sample was then lowered (1°C/min) to that desired for the particular run. The longer DNA samples were brought directly to the temperature of the run from ambient temperature (∼24 °C). Runs consisted of allowing the sample to equilibrate (between denatured single and natured double strands) for approximately 40 minutes at the given temperature and then inserting and retracting a shutter to cycle the deuterium light on and off through the sample for different time periods while continuously monitoring the wavelength dependent extinction, Eq. (1), during light on periods.

## Results

Figure 1 shows a typical extinction spectrum using the deuterium light source for (a) 48 bp synthetic DNA and (b) yeast DNA, at three different temperatures. The extinction peak at 260 nm is due predominantly to absorption on the nucleic acid bases but also includes a small amount of Rayleigh and Mie scattering. Hyperchromism is observable at the absorption peak at 260 nm due to temperature denaturing. The inset shows an amplified view of the longer wavelength region for the three different temperatures. The extinction in this region is due to Mie scattering dominating in the short wavelength region (∼200 to 350 nm) and Rayleigh scattering dominating in the long wavelength (>350 nm) region for the 25 bp and 48 bp synthetic DNA (physical size ∼ 2 × 17 nm). For the much longer yeast and salmon sperm DNA, Mie scattering is predominant in this region (Fig. 1(b) inset).

**Fig. 1.**
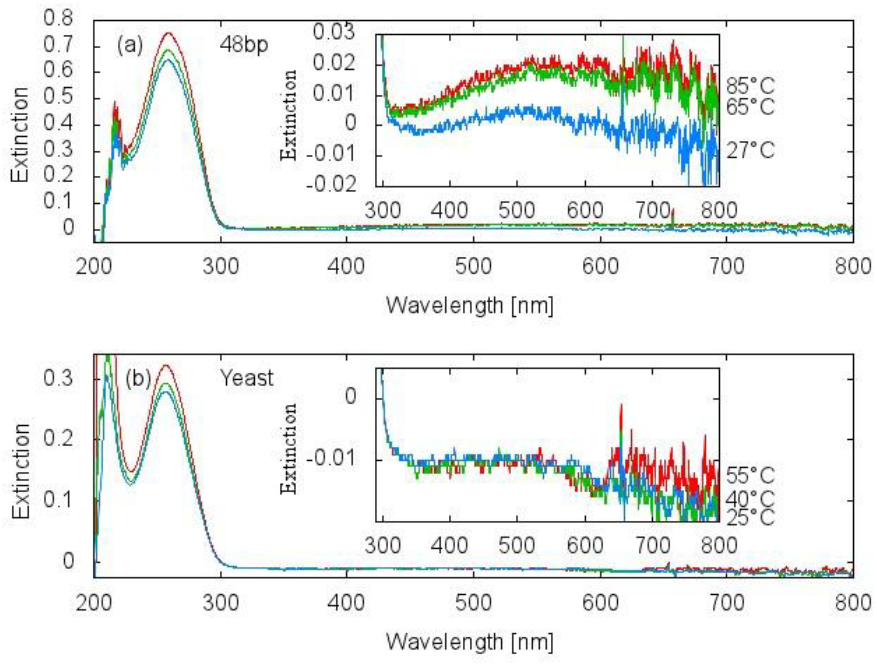
Extinction Spectra.

a) 48 bp synthetic DNA. b) 100 kbp (average size) yeast DNA. Hyperchromism is observable at 260 nm in both samples. The insets show an expansion of the scattering regions at three different temperatures.

Scattering is a complicated function of the number of scattering centers, the ratio of the indices of refraction DNA/buffer, and shape of the scattering centers^21^. However, for the small 48 bp (inset, figure 1(a)) and 25 bp (not shown) DNA the scattering increases with denaturing (correlated with temperature), while the scattering is constant or only slightly increasing with denaturing for the large yeast (inset, figure 1(b)) and salmon sperm DNA (not shown). This result is interpreted as being due to complete denaturing for at least some of the small synthetic DNA (even at low temperature) where the separated strands act as separate scattering centers, but only partial denaturing for the large DNA, where the still united strands act as single scattering centers. Scattering is thus an additional (to hyperchromicity) sensitive indicator of denaturing for the small synthetic DNA, while it has little utility for the long DNA, except at high temperature where complete denaturing also occurs. For the short synthetic DNA, therefore, a sensitive measure of photon induced denaturing was obtained by integrating the extinction over wavelength from the absorption peak to up to 495 nm.

Figures 2(a), 3(a), 4(a) and 5(a) plot the extinction, Eqn. (1), over the duration of the run, integrated over 245-295 nm and 255-275 nm, for the yeast and salmon sperm DNA, and integrated over wavelengths 245-495 nm for the short synthetic DNA samples respectively. The periods of deuterium light cycling on and off are noted on the graphs by up and down arrows respectively, at the given fixed temperature. It can be observed that in all cases, as a result of UV-C light induced denaturing, the extinction rises during the light on periods, due to absorption hyperchromicity, plus, for the short DNA, increased Rayleigh or Mie scattering.

**Fig. 2.**
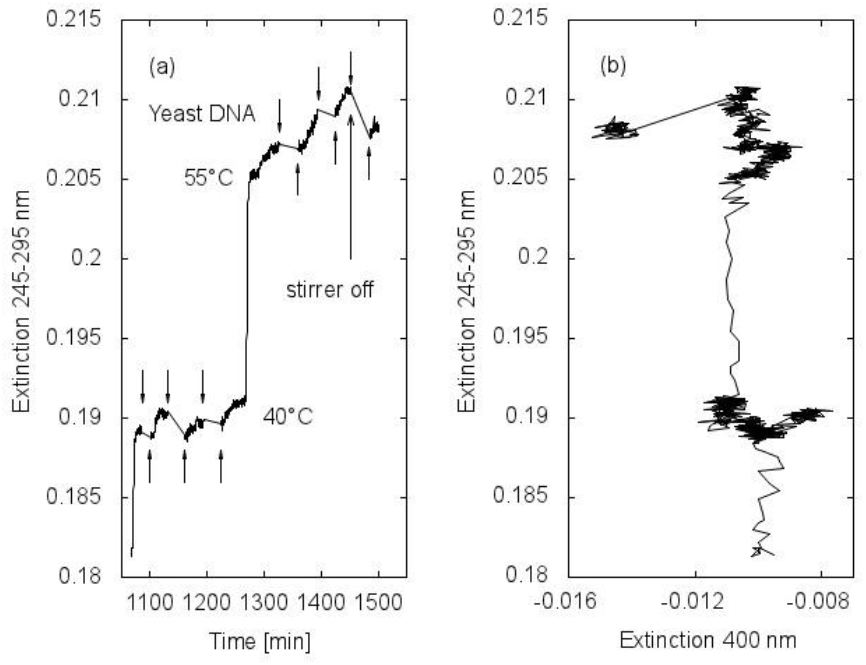
Absorption and Absorption-Scattering Correlation for Yeast DNA.

a) Extinction (mainly absorption) as a function of time in minutes. Deuterium light on an off changes are noted on the graphs by up and down arrows respectively. A change in the stirrer condtion is noted on the graph for yeast DNA. b) Correlation between extinction (mainly absorption) and Rayleigh or Mie scattering at 400 nm.

**Fig. 3.**
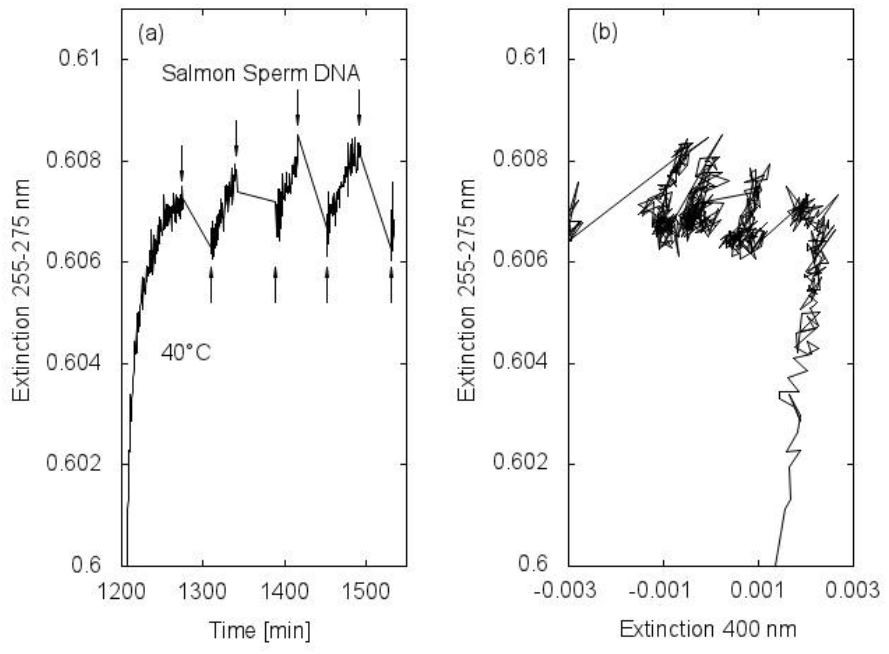
Absorption and Absorption-Scattering Correlation for Salmon Sperm DNA.

See Fig.2 for an explanation.

**Fig. 4.**
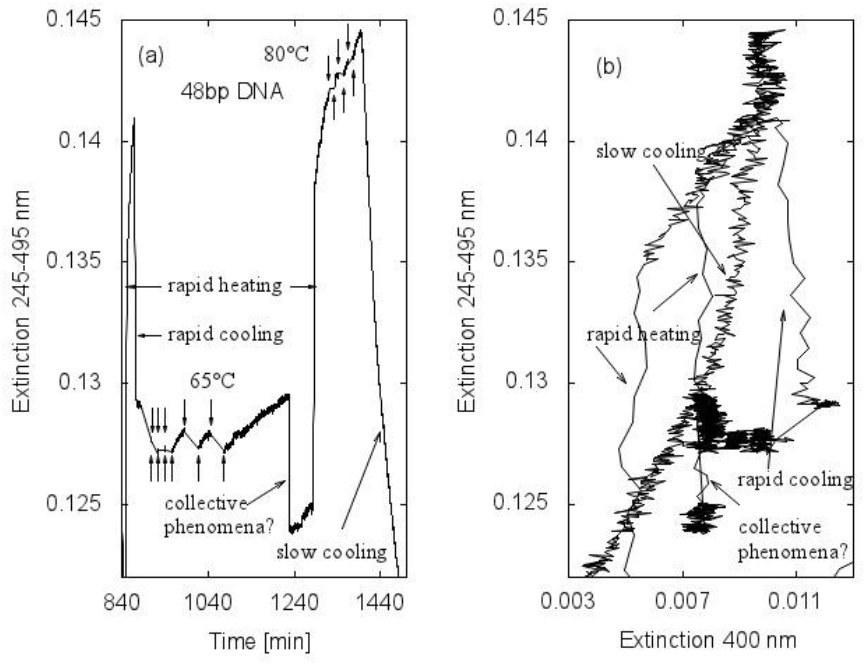
Absorption and Absorption-Scattering Correlation for 48 bp Synthetic DNA.

See Fig.2 for an explanation.

**Fig. 5.**
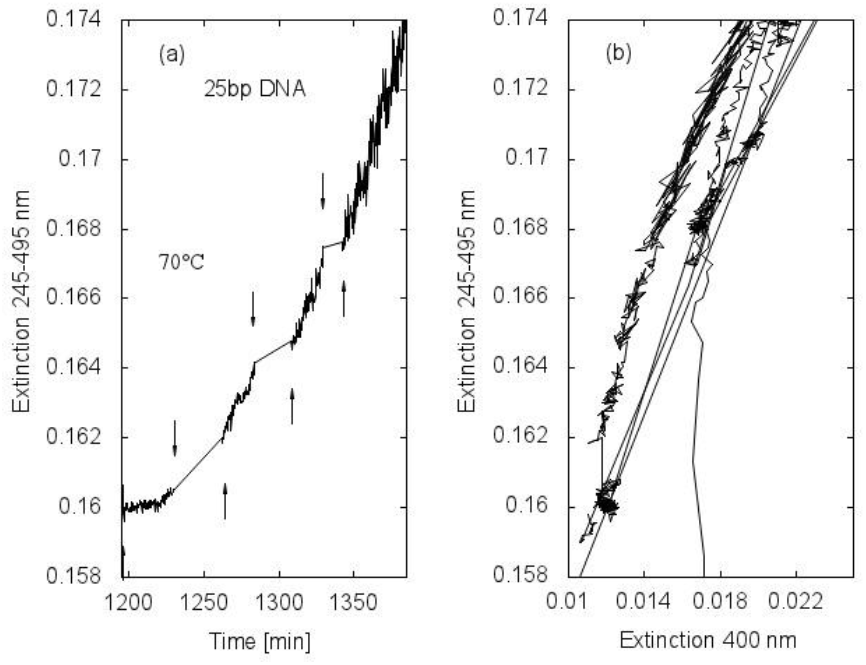
Absorption and Absorption-Scattering Correlation for 25 bp Synthetic DNA.

See Fig.2 for an explanation.

Figures 2(b), 3(b), 4(b) and 5(b) show the correlation spectrum of the extinction peak versus only scattering at 400 nm for the different DNA samples over the duration of the runs. From these figures, particularly 4(b) for 48 bp DNA, it can be deduced that while the heating or cooling is slow (0.2 °C/min) there is strong correlation between the absorption and scattering, but little correlation is seen during rapid heating or cooling. This was interpreted as being due to the fact that heat separation of the DNA bases at hydrogen bonding sites allows arbitrary orientation of the bases around the axis of the DNA giving rise to hyperchromicity, but this does not imply the physical separation of the complementary strands to an extent (» one wavelength) that would make them separate Rayleigh or Mie scattering centers. Thus, the increase in scattering lags behind the increase in absorption due to a finite diffusion time constant since it requires substantial separation of the liberated strands in order for them to act as separate scattering centers. The time constant is dependent on factors such as the salt concentration and the stirrer setting.

Figures 2(b) and 3(b) indicate that Mie scattering increases only slightly with absorption for the long yeast and salmon sperm DNA (∼100 kbp) as compared to the synthetic 25 bp DNA (Fig. 5(b)). This was interpreted as being due to the fact that photon dissipation induces only partial denaturing in these long segments while usually complete denaturing of the short synthetic DNA, particularly for the 25 bp DNA. Partial denaturing does not allow the complementary segments to separate sufficiently to act as independent Mie scattering centers.

Renaturing of completely separated DNA proceeds via chance encounter of a few complementary base pairs and then rapid re-zipping of the remaining pairs. It is thus second order in the concentration but there is also a volume or steric effect since for longer DNA the folding of single strands implies that some base pairs are inaccessible to first encounter. Renaturing is thus a diffusion plus steric limited process but has some peculiarities that depend on the nucleotide complexity of the strand, the ionic nature of the solvent, and also the viscosity of the solvent^22^. Renaturing is readily observed as a decrease in absorption after the end of the light off period in the photon induced partial denaturing of yeast and salmon sperm DNA (Figs. 2(a) and 3(a)) and to some extent in the 48 bp DNA (Fig. 4(a)) but not observed, as expected, for the completely denatured short 25 bp synthetic DNA, where, on the contrary, it appears that denaturing continues during the light off periods; probably due to heat denaturing once UV-C light induced partial denaturing has reduced the melting temperature of the surviving double strand segments (see Fig. 5(a)).

“Collective phenomena” marked on the 48 bp DNA graph (Fig. 4(a)) refers to one of a number of unexpected, and still unexplained, sudden decreases (within 30 seconds) of the extinction, found usually after a prolonged light on period for both the synthetic 25 bp (not shown) and 48 bp DNA, but not found for the long DNA. Our initial suspicion is that this may be attributed to sudden collective form changes from the canonical B-stacking geometry, for example, a B-Z transformation, perhaps a result of UV light ionization of DNA reducing the electrostatic repulsion of the phosphate residues^23^. The fact that there is no change in the scattering observed during the sudden drop in extinction (see Fig. 4(b)) indicates that this phenomena is probably not agglomeration of the DNA strands.

A quantitative measure of the UV-C light induced denaturing can be obtained by comparing the increase in hyperchromicity obtained during the light on periods with the maximum temperature induced hyperchromicity of completely denatured DNA at 90°C, assuming a linear relation between hyperchromicity and denaturing. For the salmon sperm DNA in purified water at 40 °C (at the beginning of the denaturing curve), a half hour light on period is sufficient to increase the extinction in the 255-275 nm region by 0.002 units (see Fig. 3(a)), which corresponds to 1.7% of the maximum hyperchromicity at 90°C, implying a rate of UV-C light induced denaturing of approximately 3.4% per hour. A similar value was found for yeast DNA. An analogous calculation for the 48 bp DNA in PBS buffer (see Fig. 4(a)) shows that at 65 °C (at the beginning of denaturing) the rate of UVC-light induced denaturing is approximately 4.6% per hour. For the 25 bp DNA in PBS buffer (see Fig. 5(a)) at 70 °C (somewhat into the denaturing curve) the UV-C light induced denaturing is approximately 20.1% per hour.

In order to verify that the increase in extinction observed for all samples during the light on periods was in fact due to UV-C light induced denaturing and not to some other mechanical, optical, electronic, or temperature artifact, we carried out an identical experiment with 48 bp DNA replacing the deuterium UV-C lamp with the control halogen-tungsten visible lamp. In this case (data not shown), no increase in the extinction (which could only be observable in scattering) was found during the light on sample periods, suggesting that the observed increase in extinction with the deuterium light is, indeed, due to DNA denaturing through UV-C photon dissipation.

Finally, it is pertinent to compare our experimental conditions of light, temperature and DNA concentrations to those likely existent during the Archean at the beginning of life on Earth (∼3.8 Ga). Sagan^9^ has calculated an integrated UV-C flux during the Archean over the 240-270 nm region where DNA absorbs strongly (see Fig. 1) of 3.3 W/m^2^ while we have estimated a somewhat larger flux of 4.3 W/m^2 [1]^. Our light on sample was estimated to have a flux over this same wavelength range of 2.9 W/m^2^. However, the beam volume on sample was only 0.107 cm^3^ while the total volume of the sample was 1.0 cm^3^ and uniform mixing of the DNA sample due to the magnetic stirrer ensured homogeneous passage of the DNA through the whole volume, giving an effective UV-C light flux of only 0.31 W/m^2^ during light-on periods, about 1/10 of what it may have been in the Archean.

Isotopic geologic data suggest that at 3.8 Ga the Earth was kept warm by CO_2_ and CH_4_, maintaining average surface temperatures around 80°C^24,25^ and falling to 70 ± 15 °C at 3.5−3.2 Ga^26^. Of course, polar regions would have been colder and equatorial regions warmer. The data presented here shows that at these high temperatures UV-C light induced denaturing is significant and larger than at colder temperatures.

Miller^27^ has determined adenine concentrations of 15 μM in the prebiotic soup using calculations of photochemical production rates of prebiotic organic molecules determined by Stribling and Miller^28^. Although these estimates are considered overly optimistic by some, it is noted that Miller was not aware of non-equilibrium thermodynamic routes to nucleotide proliferation based on photon dissipation^11^, for example, the recently discovered route to pyrimidine ribonucleotide production utilizing UV-C light^29^ or the purine production in UV-C irradiated formamide solutions^30^, nor of the existence of an organically enriched sea surface skin layer^31,32^, so Miller’s determinations of nucleotide concentrations at the beginnings of life may, in fact, be overly conservative. Adenine concentrations in our sample DNA were of the order of 50 μM.

The above comparisons, particularly our lower on-sample UV-C light flux relative to estimates of the flux at Earth’s surface during the Archean, suggest that our determined UV-C induced denaturing rates should probably be taken as a conservative lower limit to rates that could have occurred on the Archean sea surface at the beginning of life.

## Discussion

Through an analysis of extinction (absorption plus scattering) at UV-C wavelengths and only scattering at longer wavelengths, and considering the hyperchromicity and Rayleigh and Mie scattering characteristics of DNA, we have demonstrated that absorption and dissipation of UV-C light in the range of 230 to 290 nm denatures DNA in a reversible and apparently benign manner. Analysis of the scattering component suggests complete denaturing for short 25 bp synthetic DNA while partial denaturing for long salmon sperm, yeast DNA, and a mixture of the two for 48 bp DNA. Suspicion of UV light induced denaturing of DNA has been given previously in a different context^33^. Our results clearly indicate that UV-C light denatures DNA and we have quantified this effect under conditions that may be considered conservative with respect to those expected at the beginnings of life in the Archean.

Since RNA has similar optical and chemical properties as DNA, it is plausible that such replication, homochirality, and information acquisition mechanisms driven by photon dissipation would have acted similarly over RNA. We are presently conducting experiments to verify this. If the latter is borne out, then DNA and RNA may have been contemporaries in the primordial soup and evolved with other fundamental UV-C absorbing molecules independently towards increased photon dissipation before finally forming a symbiosis which augmented further still this dissipation. Evidence for a stereochemical era in which amino acids and other UV-C and visible absorbing fundamental molecules of life had chemical affinity to both DNA and RNA^1,4^ supports this assertion. This would have allowed continued replication and proliferation, and thus increases in photon dissipation, despite the cooling of the ocean surface and the eventual attenuation of the UV-C light brought on by the accumulation of oxygen and the delegation of UV-C dissipation to life derived ozone in the upper atmosphere.

Regarding the second part of the UVTAR mechanism, that of enzyme-less extension during dark and cooler, overnight, periods, or during cooler early morning periods, there exists experimental data obtained using chemical activation indicating that long random sequence linear DNA oligonucleotides can be template synthesized from a random pool of short oligonucleotides with high fidelity (the fidelity increasing with temperature)^34^. In the UVTAR scenario, chemical activation would be replaced by UV-C activation.

The association of growth, replication, and evolution with dissipation is a necessary thermodynamic prerequisite of any irreversible biotic or abiotic process and in this context our result could reconcile “replication first” with “metabolism first” scenarios for the origin of life. Photon dissipation is certainly the most primitive and extensive of all “metabolisms”. The results presented here also lend plausibility to the proposal of the acquisition of homochirality and information content in DNA through dissipation^7,8,17^. The thermodynamic dissipation theory for the origin of life^7,8^ suggests that all biotic, and coupled biotic-abiotic, evolution has been driven by increases in the entropy production of the biosphere, mostly through increasing the global solar photon dissipation rate.

## Acknowledgements

The authors are grateful for the DNA samples provided by Laura Ongay and Yolanda Camacho of the Institute of Cellular Physiology at the UNAM and to the financial support of DGAPA-UNAM project number IN-103113 and to CONACyT for financial support to N.S.P.

